# Transcriptome data analysis reveals correlation between developmentally altered gene expression and offspring phenotype in epigenetic inheritance

**DOI:** 10.1101/048256

**Authors:** Abhay Sharma

## Abstract

A recent study investigated sperm mediated inheritance of diet induced metabolic phenotype, reported underlying regulation of MERVL targets and ribosomal protein genes in embryos, and suggested that the altered regulation observed may cause placentation defects which can secondarily result in abnormal metabolism. An analysis of available transcriptomic data however reveals a direct link between the developmentally altered genes and the offspring phenotype.

---

Accumulating evidence suggests that parental exposure of certain environmental factors can cause altered phenotypes in subsequent generation(s). Mechanisms underlying this unconventional mode of transmission however remain elusive. Importantly, Sharma *et al.* (*1*) demonstrated sperm mediated transfer of phenotypic information in a mouse model of paternal Low Protein diet induced offspring metabolic perturbation. Upon finding altered small RNA population including increased levels of tRF-Gly-GCC in Low Protein sperm, the authors examined the transcriptomic effect of antisense oligos targeting the tRF on ESCs and embryos, and obtained evidence suggesting tRF-Gly-GCC regulation of MERVL targets. This evidence was further supported by RNA-seq analysis of embryos cultured to various stages of development following IVF with Control or Low Protein sperm or ICSI, and embryos generated through zygotic injection of sperm small RNA population or synthetic tRF-Gly-GCC oligos. Besides, Sharma *et al.* also found that altered transcripts in Low Protein sperm IVF embryos are enriched for ribosomal protein genes. The authors, instead of examining if a correlation exists between embryonic gene expression changes and offspring phenotype, explained these findings by speculating that MERVL target and ribosomal protein gene regulation may lead to altered placentation which in turn can cause downstream effects on metabolism. The present analysis of Sharma *et al.'s* gene expression data however directly links gene expression alterations in embryos with offspring phenotype, thus providing a new interpretation of the reported findings.

First, a gene ontology analysis of the previously identified MERVL targets (*2*) that formed the basis of Sharma *et al.'s* evidence itself shows that these targets overrepresent various processes relevant to the offspring phenotype described for the mouse model (*1*, *3*) (Fig. 1A). Second, the genes identified by Sharma *et al.* as differentially expressed in embryos generated following zygotic microinjection of antisense tRF-Gly-GCC oligos (table S6 of Sharma *et al*.) show similar enrichment (Fig. 1B). Third, the genes showing expression change in Low Protein sperm IVF embryos (table S7 of Sharma *et al*.) at 2-fold cut-off, a criterion used by the authors for analyzing differential expression in embryos (Fig. 4E of Sharma *et al*.), enrich several gene ontology categories relevant in epigenetic inheritance of metabolic phenotype (Fig. 2A). Fourth, similar enrichment of processes is observed in the genes showing 2-fold expression change in embryos generated through Low Protein sperm ICSI, and zygotic injection of sperm small RNA population or synthetic tRF-Gly-GCC oligos (table S8 of Sharma *et al*.) (Fig. 2B). Cumulatively, this reanalysis establishes embryonic gene expression-offspring phenotype correlation.

**Fig. 1.**
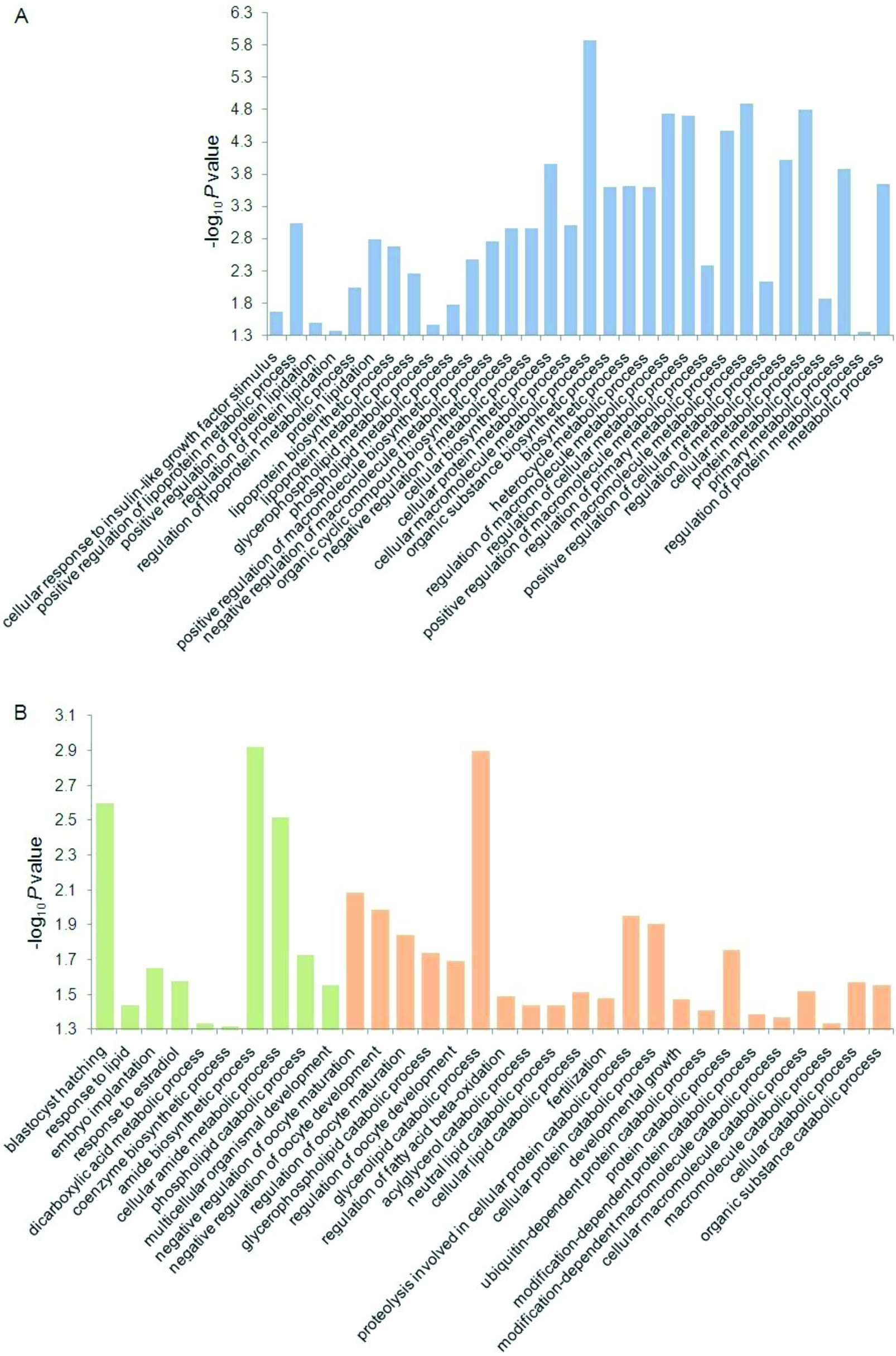
Enriched processes in (A) MERVL targets, and (B) tRF-Gly-GCC inhibition induced differentially expressed genes in embryos. Green and orange bars represent down‐ and up‐ regulated genes, in that order. Nominal significance *P* values (y axis, ‐log_10_) for enrichment are shown. Gene ontology tool (*5*) was used for enrichment analysis. Not all enriched processes are shown.

**Fig. 2.**
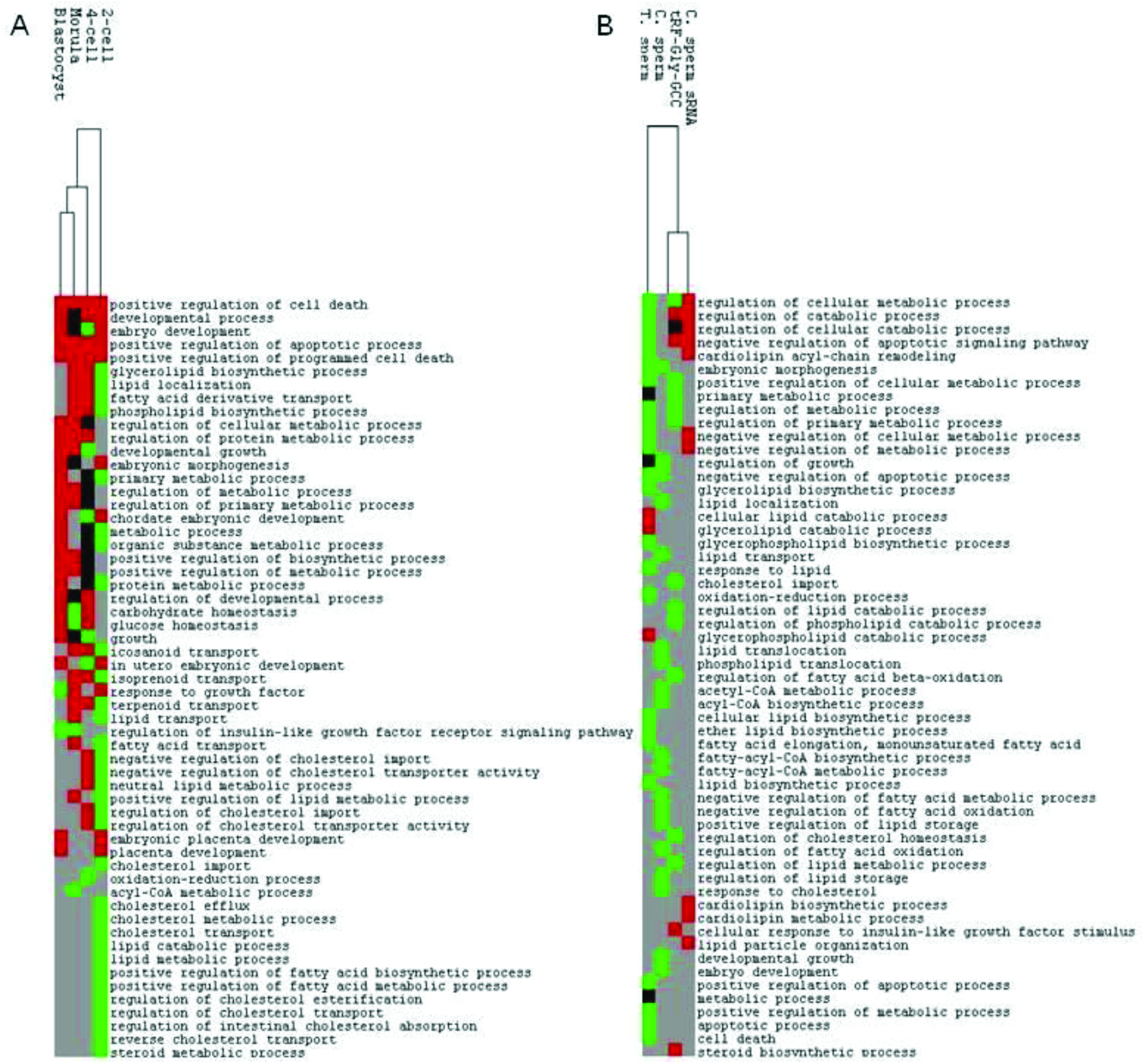
Heatmap clustering of enriched processes in differentially expressed genes in (A) Low Protein sperm IVF generated embryos at various stages, and (B) 2-cell stage embryos obtained using zygotic sperm small RNA or synthetic tRF-Gly-GCC injection, and T. sperm or C. sperm ICSI. Red, green, and black represent enriched processes in up‐ and down‐ regulated genes, and both, in that order. Grey represents no enrichment. City-block distance was used as similarity metric and average linkage as hierarchical clustering method. The data was organized and analyzed using Cluster 3.0 (*6*), and the results graphically represnted using Java TreeView 1.1.6r2 (*7*). Other details as in Fig. 1.

Like Sharma *et al.'s* article, a recently published paper separately reported sperm tRF mediated inheritance of diet induced metabolic disorder (*4*). In this study, RNA-seq analysis of 8-cell and blastocyst stage embryos generated through zygotic injection of High Fat sperm tRFs identified differentially expressed genes that were enriched for gene ontology processes related to metabolic regulation, besides others. It was suggested that these embryonic transcriptional changes may lead to reprogrammed gene expression and result in offspring phenotype. The present reanalysis is consistent with this hypothesis. Besides tRFs, these studies, as also others reporting recently epigenetic inheritance of metabolic disorders through the male line (*8*, *9*), also found altered sperm levels of other small noncoding RNAs including let-7 miRNAs. Interestingly, a role of miRNAs, specifically let-7 species, in epigenetic inheritance was predicted previously on the basis of a bioinformatic analysis *(*10*)* that tested the proposal that small RNAs mediate the transmission of environmental effects across generations through gene networks (*11*-*13*). Emerging evidence (*1*, *4*, *8*, *9*) seem consistent with this model.

## Acknowledgements

The laboratory of AS is supported by the research grant BSC0122 of the Council of Scientific and Industrial Research, India.

